# Reward and punishment differentially shape basketball free-throw learning

**DOI:** 10.64898/2026.04.28.721312

**Authors:** Charalambos Papaxanthis, Lionel Crognier, Elias Pibarot, Jérémie Gaveau, Célia Ruffino, Pierre Vassiliadis

## Abstract

Motor learning is shaped by motivational context: punishment can accelerate initial learning, whereas reward enhances memory retention. Yet it remains unclear whether the dissociable effects of reward and punishment observed in laboratory tasks generalize to complex real-world skills. Here, we tested this idea using a naturalistic motor task—basketball free-throw shooting. Sixty-eight participants trained under four motivational contexts that differed only in how points were awarded for each pair of consecutive shots: control (standard scoring), reward (bonus points for two consecutive successful shots), punishment (penalty for two consecutive missed shots), or mixed (both bonus and penalty). Performance was assessed before training, immediately after, and 1 and 3 days later. Punishment and mixed schedules significantly improved early acquisition, resulting in higher accuracy immediately after training compared to the control and reward conditions. This advantage emerged during the first training block, indicating a rapid motivational influence on performance. In contrast, reward selectively enhanced offline consolidation: three days after training, the reward group showed the largest gains in accuracy, outperforming both the control and punishment groups. The mixed schedule produced quick early gains similar to punishment, but achieved smaller long-term improvements than reward. Consistent with these findings, individual punishment sensitivity was associated with gains in acquisition, while reward sensitivity correlated with offline improvements. Together, these findings demonstrate dissociable effects of motivational valence on the acquisition and consolidation of a complex real-world motor skill. More generally, they position motivational interventions as simple and cost-effective strategies to enhance rehabilitation and sports training.

## Introduction

Motor learning, the ability to refine movement kinematics through practice, is strongly shaped by the motivational context in which training occurs. Over two decades of research have suggested that reward and punishment can differentially influence components of motor learning in various laboratory tasks, including sensorimotor adaptation^1–3^, sequence learning^4,5^, and force modulation^6–9^.

This body of work shows that reward can reliably enhance the retention of motor memories (^1,5,9–11^; but see also ^12^ for a null result), with benefits lasting up to 30 days after training^6^. In contrast, several motor adaptation studies report that punishment preferentially facilitates initial, rapid online learning^1,13,14^. These findings suggest that motivational feedback can differentially shape distinct phases of motor learning. This raises the possibility that motivational interventions may offer efficient, low-cost avenues to enhance motor learning in real-world contexts such as motor rehabilitation^15,16^ and sport performance^17,18^. Yet it remains unclear whether and how these benefits extend to more complex, real-world skills.

Providing reward or punishment after movement can modulate learning via two partly dissociable mechanisms. First, delivering explicit outcomes (e.g., success vs. failure) supplies knowledge of performance that can, on its own, drive trial-by-trial motor adjustments essential for learning^2,7,18–23^. Second, the motivational context in which a skill is acquired—manipulated by increasing the value of good performance (e.g., via monetary incentives) while keeping knowledge of performance matched across conditions—can also profoundly impact learning^5,7,24^. A complementary way to modulate motivation is to change outcome valence, i.e., whether good or bad performance yields positive or negative reinforcement (e.g., points gained or lost^1,6,10,14^).

Manipulating outcome valence through points may constitute a particularly attractive motivational intervention for more complex, real-world skills, especially in rehabilitation and sports training. First, it avoids monetary incentives, which can be unrealistic for day-to-day practice and may raise ethical concerns—particularly for patients and young practitioners. Second, the dissociable effects of reward and punishment observed in laboratory tasks may be harnessed to target different learning processes depending on training needs. Third, combining reward and punishment in a “mixed” schedule to potentially capture the benefits of both. Accordingly, in this study, we tested whether punishment and reward effects generalize to a real-world task, basketball free-throwing skill, by preferentially augmenting online acquisition and offline consolidation, respectively, and whether their combination yielded additional gains. Crucially, the reward, punishment, and mixed conditions only differed in how points were computed (bonuses vs. penalties; see below), with no monetary incentives. This setup allows us to evaluate the applicability of this approach in a realistic scenario compatible with daily rehabilitation and sport-coaching practice.

## Results

Sixty-eight healthy adults were randomly assigned to four groups (n = 17 each) and completed a basketball free-throw task across three sessions. In session 1, between Pre-training (Pre) and Post-training (Post) assessments (10 throws each), participants trained on the task for 120 trials (10 blocks of 12 trials), allowing us to assess motor skill acquisition (Fig. 1A). Sessions 2 and 3, each including 10 throws, were conducted 1 day (Post_+1d_) and 3 days (Post_+3d_) after session 1 to assess offline skill consolidation (see Fig.1A). The free throws were performed 2 by 2, mimicking real basketball practice, with brief rest periods between shots.

**Figure 1.**
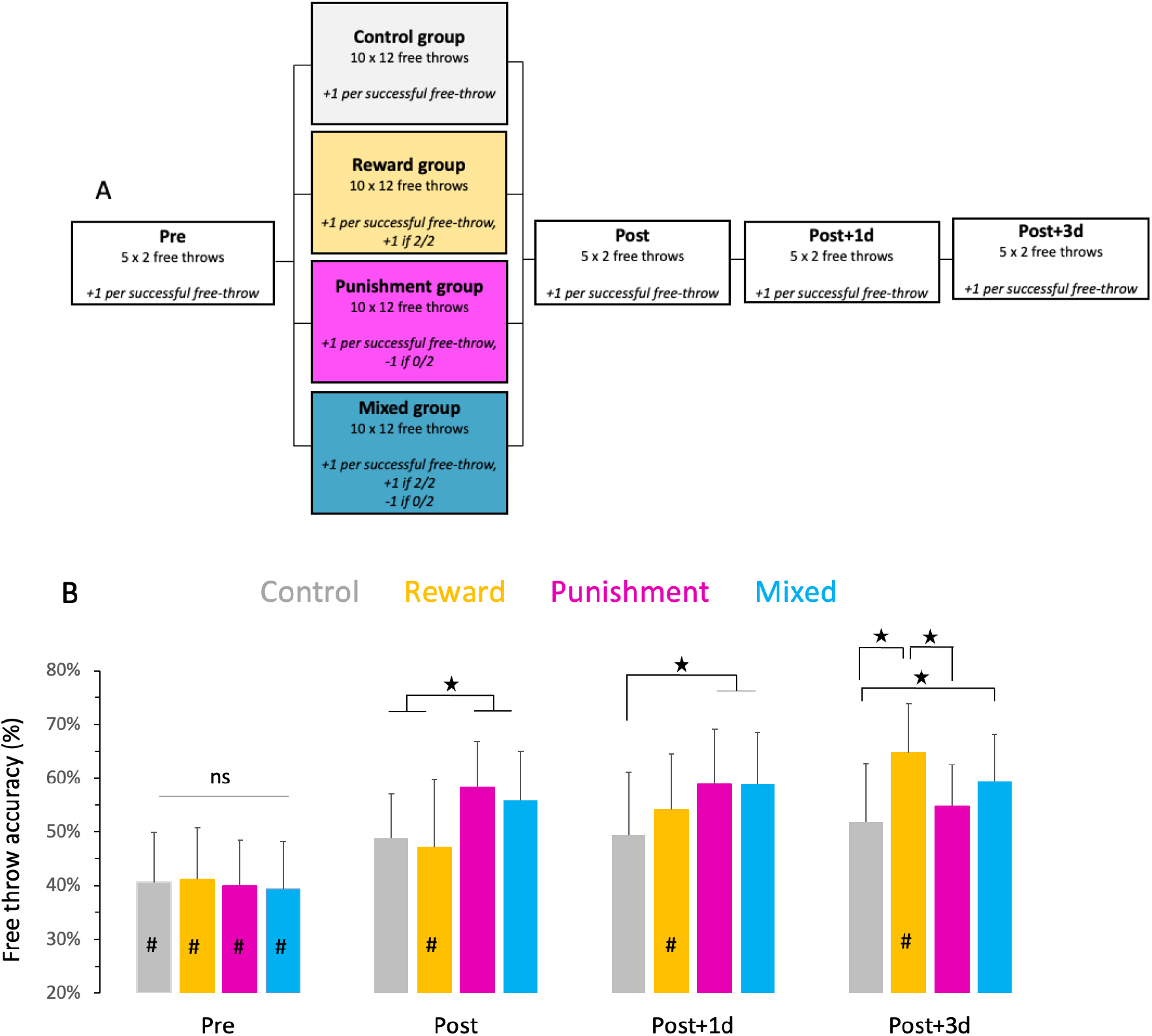
**(A) Experimental protocol.** In the Pre-training test (Pre), all participants completed 10 free throws (in blocks of 2 consecutive shots). All had full sensory feedback during their free throws, but no verbal feedback or instruction from the experimenter. In Pre, all participants received 1 point for each successful free throw, according to the conventional scoring procedure. During training, participants were divided into four groups (see the different colours) and performed 120 free throws (i.e., 10 rounds of 12 free throws, with 30 seconds of rest between series). Within each round, participants completed the free throw in blocks of two consecutive shots. Throughout training, all groups had full sensory feedback and received one point if one of the two free throws was successful, but they varied in the number of points awarded if both free throws were missed (0, 0, -1, -1 for the control, reward, punishment, and mixed groups, respectively) or if both free throws were successful (2, 3, 2, 3 for the control, reward, punishment, and mixed groups, respectively). Immediately (Post), one day (Post_+1d_), and three days (Post_+3d_) after training, all participants performed ten free throws as in the Pre, using the conventional scoring. **(B) Average (% + SD, Standard Deviation) free-throw accuracy for the four groups across the four test sessions**. Each training group is represented with a different colour. Stars (*) indicate significant differences between groups within each test session. Inside each group’s histogram, hashtags (#) indicate a significant difference between that histogram and the others of the same colour.

During training, we manipulated participants’ motivation by varying how points were counted across groups. In the *control* group, scoring was exactly as in standard basketball (0, 1, or 2 points for 0, 1 or 2 successful shots, respectively). In the *reward* group, the difference from the control group was that participants could earn a bonus point for scoring two consecutive shots in a block (0, 1, or 3 points). In the *punishment* group, the difference from the control group was that participants could lose a point if neither of the 2 consecutive free-throws were successful (-1, 1, or 2 points). The *mixed* group combined features of the reward and punishment groups (-1, 1, or 3 points). This design allowed us to examine how different motivational contexts influence the acquisition and consolidation of a real-world motor skill. Proportions of successful throws were analysed using non-parametric tests, focusing first on between-group differences in overall performance, then on acquisition and offline memory consolidation.

### Reward and punishment increase long- and short-term free-throw learning, respectively

As shown on Fig. 1B, there was no initial difference in free-throw accuracy between groups, as measured at Pre (Kruskal-Wallis ANOVA, H=0.60, p=0.90, n^2^=0.04). Mean within-group accuracy ranged from 39% to 41%, with an overall mean of 40% ± 9% (n = 68, Fig. 1B).

Free-throw acquisition at Post significantly differed between groups (Kruskal-Wallis ANOVA, H=13.06, p=0.005, n^2^=0.16). The punishment and mixed groups, although not different from each other (p=0.58, effect-size r=0.09), improved their accuracy significantly more than the control and reward training groups (in all, p<0.03, r>0.37). Control and reward groups did not differ at Post (p=0.64, r=0.08). This result suggests that the penalty component, present in both the punishment and mixed groups, enhanced online learning in the first session, whereas the bonus component had no effect initially.

To further characterise the initial benefit of the penalty, we evaluated the timing of its emergence during training. The penalty-related improvement in free-throw accuracy emerged very early, with a significant difference (Kruskal-Wallis ANOVA, H=9.18, p=0.027, n^2^=0.09) already detectable in the first training block and persisting across all blocks. The gains between the Pre and the first block were 12% for the control group, 8.5% for the reward group, 19% for the punishment group, and 18% for the mixed group. Mann-Whitney U tests showed that the punishment and mixed groups, although not significantly different from each other (p=0.88, r=0.02), improved their accuracy significantly more than the reward training group (all, p<0.02, r>0.40) and nearly significantly more than the control group (p=0.05, r=0.33 and p=0.08, r=0.29 for punishment and mixed groups, respectively). Control and reward training groups did not differ (p=0.22, r=0.21). This effect suggests that the benefit of a penalty on skill acquisition is associated with an early boost in performance.

A significant difference among groups was also detected at Post_+1d_ (H=9.30, p=0.02, n^2^=0.10). Again, while the punishment and mixed groups did not differ (p=0.58, r=0.03), both had higher accuracy than the control group (for both comparisons, p=0.01 and r=0.44, respectively). The reward group, which improved its accuracy between Pre and Post_+1d_, did not differ statistically from any other group (in all comparisons, p > 0.12, r > 0.42).

Finally, tree-throw performance at Post_+3d_ was also significantly different between groups (Kruskal-Wallis ANOVA, H=13.55, p=0.003, n^2^=0.16, Fig. 1B). The reward and mixed groups achieved the highest accuracy, with no significant difference between them (p=0.16, r=0.22). Interestingly, the reward group reached higher accuracy than the control group (p=0.003, r=0.51) and the punishment group (p=0.004, r=0.49). This latter did not differ from the control group (p=0.32, r=0.16) nor from the mixed group (p=0.12, r=0.25). Hence, while the presence of a penalty improved early skill acquisition, reward progressively enhanced skill consolidation.

Analyses of within-group learning dynamics confirmed these effects. In the reward group, learning progressed steadily over time and reached the highest performance level of all groups at Post_+3d_. By contrast, in the other groups, performance improvements were driven mainly by early skill-acquisition gains, with progress tapering off thereafter (see Supplemental Fig. 1 and Supplementary materials for details on within-subject statistics).

### Reward enhances offline gains in free-throw accuracy

To directly test the idea that reward specifically enhances skill consolidation, we conducted an additional analysis comparing offline gains (Post_+1d_ - Post and Post_+3d_ - Post) in free-throw accuracy across the four groups (see Figure 2).

**Figure 2.**
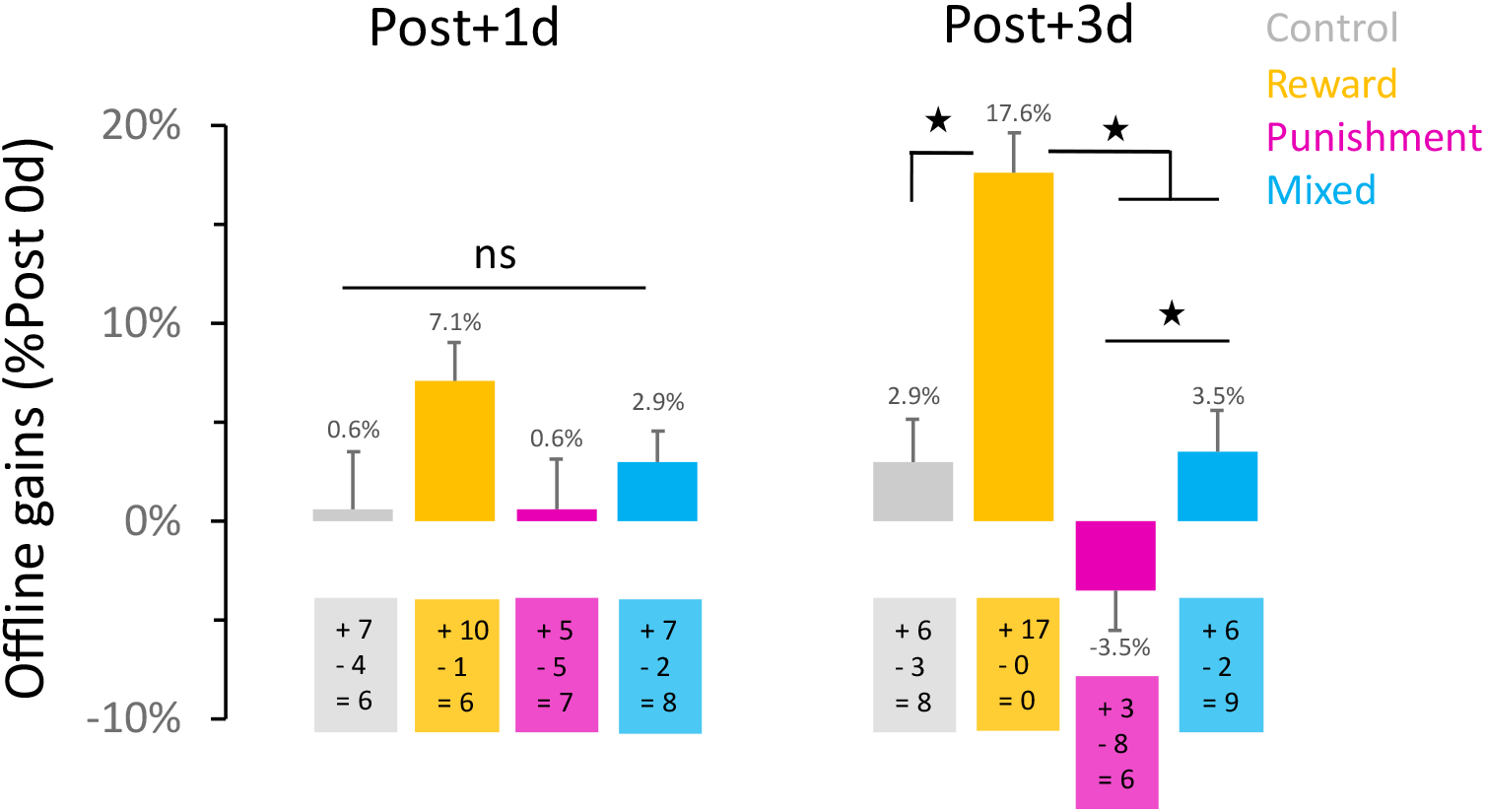
Offline gains in free-throw accuracy following training. Average improvements in free-throw accuracy (% and Standard Errors, SE) relative to Post for each group. Stars (*) indicate significant differences between groups within each test session. Each coloured box, representing each group, shows the number of participants who improved (+), decreased (-), or stabilised (=) their accuracy at Post_+1d_ and Post_+3d_ relative to Pre. Post_+1d_ (gains from Post to Post_+1d_), Post_+3d_ (gains from Post to Post_+3d_).

At Post_+1d_, free-throw gains were not different between the groups (Kruskal-Wallis ANOVA, H=4.39, p=0.22, n^2^=0.02). Note, however, that although not statistically significant, the reward group exhibited larger offline gains (+7%) than the other groups (all below +3%), with 10 participants increasing their scores from Post to Post_+1d_.

At Post_+3d_, a significant difference among groups was observed (H=30.07, p<0.0001, n^2^=0.42). The reward group showed the best consolidation (+18%), which differed significantly from the other groups (in all, p < 0.0001, r > 0.63). The punishment group’s performance decreased by 4% at Post_+3d_ compared to Post and significantly differed from the reward group (p<0.001 and r=0.78) as well as from the mixed group (p=0.03 and r=0.37). The difference between the punishment and control groups was marginal (p=0.056, r=0.33). The control and mixed groups did not differ (p=0.86 and r=0.03). Importantly, every member of the reward group (n=17) improved their accuracy from Post to Post_+3d,_ whereas only 6, 3, and 6 participants in the control, punishment, and mixed groups, respectively, showed improvement. *Fisher* tests on the number of participants who improved their accuracy revealed significant differences between the reward group and the other three groups (in all, p<0.001); the remaining comparisons did not show significant differences (all p>0.05). Overall, these results show that providing a bonus during training improves consolidation of a real-world skill, leading to better performance 3 days after training.

### Individual sensitivity to reward and punishment is associated with distinct learning dynamics

By manipulating the reward and punishment outcomes associated with training performance, the analyses above show distinct effects on the acquisition and consolidation of a motor skill. To further investigate the effects of reward and punishment, we examined individual differences in sensitivity to both using the Sensitivity to Punishment and Sensitivity to Reward Questionnaire (SPSRQ) across all participants, regardless of their training group. We then evaluated whether these individual measures were related to motor skill acquisition and consolidation.

First, we tested whether sensitivity to reward and punishment correlated with the magnitude of acquisition gains (Post – Pre). Punishment sensitivity positively correlated with acquisition gains (n=68, Kendall’s Tau *t*=0.23 and p=0.005, Fig.3A). This result was replicated when excluding the participants who did the questionnaire after training (remaining n=54, t=0.22, p=0.02, see Methods) and also when performing a multiple linear regression accounting for group identity and baseline performance (n=54, p<0.01, see Supplementary materials for more details). Crucially, reward sensitivity was not correlated with acquisition gains (across all analyses, *t-values* around 0 and p-values >0.5, Fig 3B), demonstrating the specificity of the relationship between punishment sensitivity and skill acquisition.

**Figure 3.**
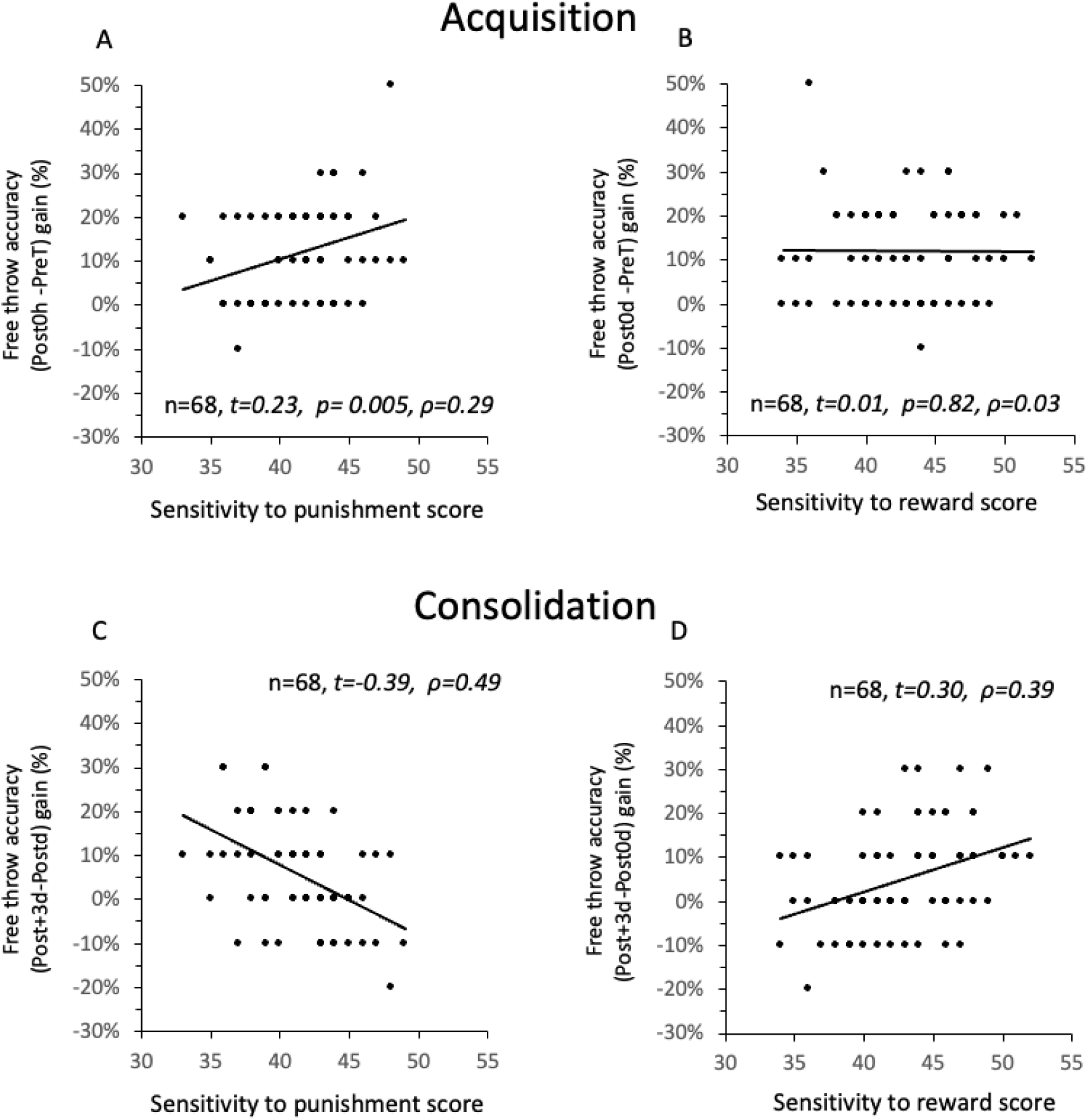
Correlational analysis. **(A)** Association between sensitivity to punishment and online free-throw accuracy gain (%, Post - Pre). **(B)** Association between sensitivity to reward and online free-throw accuracy gain (%, Post - Pre). **(C)** Association between sensitivity to punishment and offline free-throw accuracy gain (%, Post_+3d_-Post). **(D)** Association between sensitivity to reward and offline free-throw accuracy gain (%, Post_+3d_-Post). To avoid overplotting of discrete values, a small random vertical jitter was added to the y-coordinates for visualization purposes: *Y*_*plot*_=*Y* +ε, where ε ∼ *U*(−*a, a*). Correlation analyses were performed using the original data without jitter. The number of participants, Kendall’s Tau (*t*), p-values, and Spearman’s rho (ρ) are reported in the Figures.

We then used the same approach to correlate reward and punishment scores with offline gains (Post_+3d_ – Post). Punishment scores were negatively correlated (Fig. 4C) with offline gains (for n=68, *t*=-0.39; for n=54, t=-0.34; n=54, p<0.001, for multiple linear regression accounting for group identity and baseline performance see Supplementary materials), supporting that punishment training may be detrimental for long-term motor memory. Conversely, reward sensitivity was positively associated with offline gains across all analyses (Fig. 3D, n=68, *t*=0.30, p<0.001; n= 54, *t*=0.25, p=0.008; n=54, p<0.001 for multiple linear regression accounting for group identity and baseline performance in Supplementary materials). Overall, these analyses reveal that individual sensitivity to reward and punishment affects the dynamics of learning a real-world skill.

## Discussion

Across four groups of participants trained under distinct motivational contexts, we demonstrate dissociable effects of reward and punishment on the acquisition and consolidation of a basketball free-throw skill. Consistent with prior laboratory findings, we found that punishment accelerated initial skill acquisition, whereas reward markedly enhanced offline consolidation, resulting in better performance 3 days after training. The mixed schedule produced rapid early improvements, similar to punishment, that were largely maintained three days later, albeit with smaller offline gains than with reward. Finally, while online acquisition gains were correlated with punishment sensitivity, offline consolidation gains scaled with reward sensitivity. Overall, these results show that the motivational context in which training occurs, as well as individual differences in sensitivity to incentives can shape both acquisition and consolidation of a complex, real-world motor skill.

The present findings extend prior laboratory demonstrations of reward-punishment dissociations in sensorimotor adaptation^1^, sequence learning^4^, and force modulation^6^ to a complex, multi-joint skill performed under authentic sport conditions. Those effects are also consistent with recent clinical evidence that reward-based training can produce long-term, but not immediate, functional improvements in stroke patients^15^. Importantly, unlike previous work that linked performance to financial gains or losses, the present manipulation relied exclusively on points, making it directly applicable to real-world coaching settings or rehabilitation scenarios where financial rewards may be impractical or raise ethical concerns. This ecological validity is important, suggesting that these motivational mechanisms are not limited to simple laboratory paradigms or monetary incentives, but instead generalize across levels of task complexity and reward types. Beyond the consistency of the positive effect of reward in different contexts (see also ^5,7,8,16,25^), it is also worth noticing its remarkable stability within our sample, with 17/17 participants exhibiting offline gains in performance 3 days after training vs. 6/17 in the control group. Consistently, we also note that the size of the reward effect 3 days after training is large (r = 0.50 vs. the control group) which is remarkable given the cost, simplicity and duration of the intervention. In practice, the present results suggest that reward should be preferred when long-term performance is the goal. If very rapid changes in behavior are also desirable, a mixed schedule may provide a good trade-off between rapid acquisition and reasonable consolidation.

Mechanistically, the early advantage of punishment may reflect the motivational salience of potential losses. According to prospect theory^26^, losses loom larger than equivalent gains, and loss contexts can increase on-task attention and performance monitoring^28,30^, potentially facilitating rapid online adjustments. This explanation is consistent with evidence showing that punishment-induced loss aversion accelerates rapid motor correction in adaptation^1,27^ as well as sequence learning tasks^4^. This interpretation is further supported by the early emergence of the penalty benefit (first training block), consistent with a rapid, motivation-driven response rather than a slow error-based learning process. Whether this constitutes genuine motor learning or a performance-level effect is difficult to disentangle from behavioral data alone; the persistence of punishment gains at Post suggests at least partial encoding of the skill. By contrast, reward appears to be beneficial in the long run, consistent with a large body of evidence indicating that reward boosts neural mechanisms involved in consolidation of memories, including neural replay in the hippocampus^29,31,32^.

Beyond group-level effects, the present data indicate that individual motivational profiles are linked to distinct learning dynamics: sensitivity to punishment was positively associated with online acquisition gains, whereas sensitivity to reward was positively associated with offline consolidation. This pattern can be interpreted in light of evidence that higher reward sensitivity is associated with stronger ventral striatal responses during reward receipt^36^. In turn, ventral striatal engagement during rewarded training has been implicated in subsequent motor memory consolidation^33^, and recent causal work further supports a key role for the human striatum in reinforcement-related gains in retention^9^. In this framework, individuals who are more reward-sensitive may experience larger striatal reward signals during training, which could promote consolidation, whereas stronger punishment sensitivity may favor the engagement of a partially dissociable network, including the amygdala and prefrontal cortex^36,39,41^ which may be efficient for rapid online adjustments but come at the expense of offline gains. Overall, these findings suggest that individual sensitivity to reward and punishment is linked to how learners trade off rapid acquisition against longer-term consolidation, raising the possibility that such profiles could help stratify athletes or patients and tailor motivational schedules to individual needs.

This work also has some limitations. First, offline gains were computed relative to Post, as is classically done, but because the punishment and mixed groups achieved higher performance after training, we cannot rule out the possibility that some form of ceiling effect constrained offline gains in these two groups. Yet, reward-driven offline memory gains remain supported by the contrast with the control group which exhibited comparable Post performance and by the progressive emergence of a reward-related gain in Pre-normalized accuracy, significant at 3 days after training. Second, participants were intermediate learners, limiting generalization to completely novices or expert athletes; an issue for future investigation. Third, this proof-of-concept study included free-throw accuracy as the sole outcome measure. While this outcome is the most relevant with respect to basketball practice, richer movement metrics—including release angle, wrist kinematics, and multi-joint coordination obtainable via markerless motion capture^34^—will allow mechanistic decomposition of how motivational context shapes specific movement parameters in future work.

Taken together, these findings demonstrate that simple and cost-effective manipulations of outcome valence can selectively target acquisition or consolidation of a complex real-world motor skill, offering a principled framework for motivationally optimized training programs in sport and rehabilitation.

## Methods

### Participants

Sixty-eight (n = 68) young adults took part in the present study after providing informed consent. All had normal or corrected-to-normal vision and were free from neurological, physical, or musculoskeletal injuries. The task in our study was free-throw shooting in basketball. All participants were students at the Université Bourgogne Europe and had mastered the basic free-throw technique through moderate practice in the past (college or secondary school). Expert basketball players and novices who had never practiced basketball were excluded from the present study. This selection prevented a performance ceiling effect (i.e., no improvement due to high initial accuracy) or demotivation due to lack of success.

The participants were randomly assigned to four groups based on their training in free-throw shooting: the control group (n = 17, three females and fourteen males, mean age: 22.3 ± 1.2 years), the reward group (n = 17, two females and fifteen males, mean age: 21.7 ± 1.8 years), the punishment group (n = 17, seven females and eight males, mean age: 22.6 ± 0.9 years), and the mixed group, which received both reward and punishment (n = 17, three females and fourteen males, mean age: 22.7 ± 1.6 years). The regional ethics committee (CERUBFC-2025-01-15-004) approved the experimental design, which conformed to the standards set by the Declaration of Helsinki.

### Experimental device and procedure

The experiment was carried out as follows. Participants first completed the French version^35^ of the Sensitivity to Punishment and Sensitivity to Reward Questionnaire (SPSRQ). Note that 7 participants from the control group and 7 from the reward group responded to the questionnaire after the experiment. As a control, we verified that any link to learning could be reproduced after removing these 14 participants. In this questionnaire, the highest scores for punishment and reward sensitivity are 72 and 68, respectively.

Then, in the presence of only the two experimenters, they completed a standard 10-minute warm-up session that included specific basketball drills and 10 random shots, excluding free throws. The experimental protocol (see Fig. 1) included one training session and four tests: Pre-Training (Pre), immediate Post-Training (Post), one day after training (Post_+1d_), and three days after training (Post_+3d_). For all experiments, we used a standard basketball hoop in a quiet gymnasium. Men used a size 7 ball, while women used a size 6 ball.

At Pre-Training, all participants completed 10 free throws from the standard basketball distance (15 feet or 4 meters), divided into five blocks of 2 consecutive free throws to mimic real game conditions as closely as possible, with 5 seconds of rest between blocks. All participants received full sensory feedback on their free throws, but no verbal feedback or instruction was provided by the experimenter.

The training session started 5 minutes after the Pre. All participants completed 120 free throws in 12 blocks of 10, with 30 seconds of rest between blocks. The free throws within each block were performed as in the Pre, meaning 2 shots at a time with 5 seconds of rest between shots.

Participants in the *control* learning group received conventional training; that is, they tried to score on each trial and always received full sensory feedback on their performance to adjust and correct their free throws on a trial-by-trial ^37,38,40^basis. During the training, they accumulated points based on their accuracy, following basketball rules: 1 successful free throw = 1 point. They did not receive any verbal feedback, instructions, or encouragement from the experimenters.

Participants in the *reward* group received the same training as the control group, with the only difference being that they were rewarded: if both free throws within each block were successful, they earned 3 points (+1 point bonus); if only one free throw was successful, they received 1 point; and if neither was successful, they scored zero points. They thus had to try to accumulate as many extra points as possible by making both free throws.

Participants in the *punishment* group received the same training as the control group, with the only difference being that they were punished: if they failed both free throws within each block, they lost 1 point (-1 point); otherwise, they earned 1 point if only one free throw was successful or 2 points if both free throws were successful. Participants in the *mixed* group had a training session that combined elements from the other groups: full-vision, deducting 1 point if neither free throw was successful, awarding 3 points if both were successful, and giving 1 point if only one was successful.

Hence, all groups received 1 point if one of the two free-throws was successful, but they differed in the points awarded if both free throws were missed (0, 0, -1, -1 for the control, reward, punishment, and mixed groups, respectively) or if both free throws were successful (2, 3, 2, 3 for the same groups, respectively). Note that participants in the reward, punishment, and mixed groups received all the instructions before the experiment without any further verbal feedback, instructions, or encouragement from the experimenters, with the only difference between groups being the motivational context during training. Unlike previous studies that manipulated reward in motor learning^5–8^, the current study did not include monetary incentives; the only incentive was the number of points. Additionally, the groups were not competing with each other. Each group aimed to accumulate as many points as possible.

After the training session, all participants rested for 5 minutes before the immediate post-training test (Post). They also completed two additional post-training tests, 24 hours (Post_+1d_) and 72 hours (Post_+3d_). The three post-training tests were conducted under the same conditions as the Pre: a 10-minute warm-up followed by 5 blocks of 2 free throws each, with 5 seconds of rest between blocks, and without any performance-based bonus or penalty.

## Data analysis

For each test session, the number of successful free throws was recorded as follows:

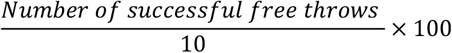

where 10 is the number of free throws in each session of pre-test (Pre) and post-tests (Post, Post_+1d_, and Post_+3d_), as well as during each block of the training session. Gains (%) in free-throw accuracy between sessions were calculated as the simple difference between them (e.g., Post – Pre).

Statistical analyses were performed using STATISTICA (8.0 version: Stat-Soft, Tulsa, OK). An a priori *G*POWER* analysis for total sample size estimation (parameters: Effect size f = 0.2; α = 0.05; power = 0.9; groups = 4; number of measurements = 4) indicated 17 participants per group (n = 68). Because the data were not normally distributed (Shapiro-Wilk test), we performed nonparametric statistical analyses. The effects of training on the four tests (Pre, Post, Post_+1d_, and Post_+3d_) within each group were tested using a Friedman ANOVA, followed by a Wilcoxon test when necessary. All the gain ratios were compared against the zero (0) value (unilateral t-tests). A Kruskal-Wallis ANOVA was used to test effects across the four groups for each test session (Pre, Post, Post_+1d_, and Post_+3d_), followed by a Mann-Whitney U test when required. The effect sizes for the Friedman ANOVA were reported as Kendall’s W (coefficient of concordance) and for Kruskal-Wallis ANOVA as n^2^. Effect sizes for Wilcoxon and Mann-Whitney U tests were reported as r 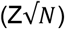. We also conducted nonparametric correlational analyses between gains in free-throw accuracy and scores related to punishment or reward. We used Kendall’s tau (τ) because many ranks were tied. The significance level for all analyses was set at 0.05.

## Supplementary materials

### 1. Within-group dynamics of skill acquisition and consolidation

As a supplementary analysis, we investigated the evolution of performance across test sessions in the different groups.

For the *control* group (grey histograms in Fig. 1B), Friedman’s ANOVA revealed a main effect of *test* (χ^2^=15.07; p=0.002; Kendall’s W=0.30). Wilcoxon tests showed a significant difference between Pre and all three post-tests (all Z>2.15, p<0.03, and r>0.52), indicating that free-throw accuracy improved immediately after training and then remained stable (Z<1.25, p>0.3, and r<0.301, for comparisons among post-tests). Supplemental Fig. 1 illustrates changes in free-throw accuracy within tests. Specifically, a +8.2% improvement (Post vs Pre, p<0.003), +0.6% (Post_+1d_ vs Post, p=0.42), and +2.4% (Post_+3d_ vs Post_+1d_, p=0.20). The total improvement in free-throw accuracy was 11.2% (p<0.001; Post_+3d_ vs Pre). Additionally, at Post_+3d_, 12 of 17 participants improved their accuracy compared to Pre, while 5 maintained it (see the grey box in suppl. Fig. 1).

**Supplemental Fig. 1.**
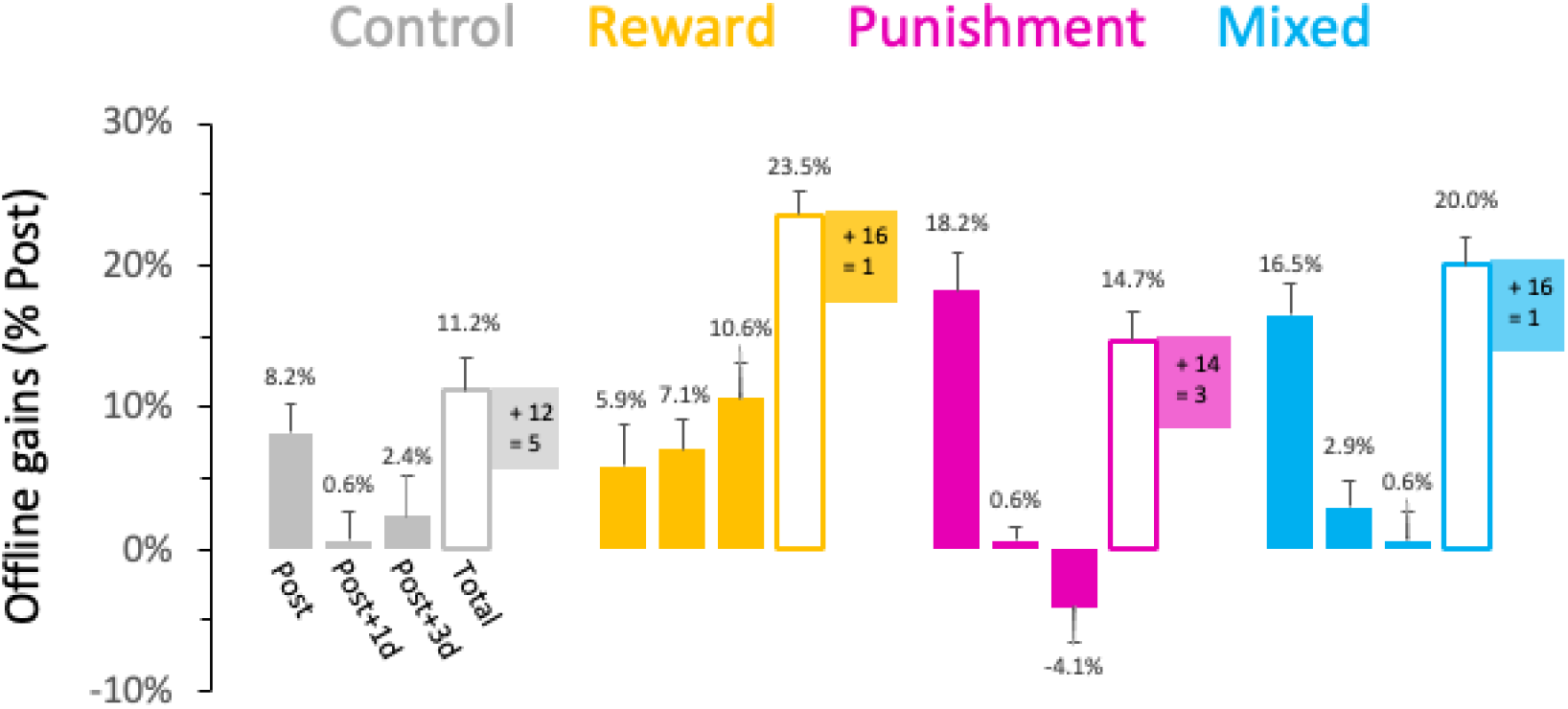
Gains in free-throw accuracy. Average improvements in free-throw accuracy (% and Standard Errors, SE) for each group from test to test. The non-filled histograms indicated the total improvement, that is, the accumulation of gains from test to test. Each coloured box, representing each group, shows the number of participants who improved (+) or stabilised (=) their accuracy at Post+3d relative to Pre (i.e., the total improvement). Post (Post – Pre), Post+_1d_ (Post+_1d_ – Post), and Post_+3d_ (Post_+3d_ – Post_+1d_).

For the *reward* group (yellow histograms), there was a main effect of *test* (χ^2^=41.59; p<0.0001; Kendall’s W=0.82). Interestingly, free-throw accuracy increased progressively from Pre to Post_+3d_, differing across all tests (all, Z>2.25, p<0.02, and r>0.55), indicating offline performance gains following the initial training session. Specifically, free-throw accuracy improvements within tests were: +5.9% (Post vs Pre, p<0.001), +7.1% (Post_+1d_ vs Post, p<0.001), and +10.6% (Post_+3d_ vs Post_+1d_, p<0.001). The total increase in free-throw accuracy was 23.5% (p<0.001; Post_+3d_ versus Pre). Notably, at Post_+3d_, 16 of 17 participants improved their accuracy compared to Pre, while 1 maintained it (see the yellow box in suppl. Fig. 1).

For the *punishment* group (magenta histograms), we observed a main effect of *test* (χ^2^=30.82; p<0.0001; Kendall’s W=0.60). Compared to Pre, free-throw accuracy improved across all post-tests (all, Z>3.30, p<0.001, and r>0.80), with no further changes (Z<1.46, p>0.1 and r<0.35 for comparisons among post-test sessions). Notably, free-throw accuracy improved remarkably immediately after training (+18.2%, Post vs Pre, p<0.001), remained stable at Post_+1d_ (+0.6%, vs Post, p=0.41), and slightly declined at Post_+3d_ (-4.1%, vs Post_+1d_, p=0.06). The overall increase in free-throw accuracy was 14.7% (p<0.001; Post_+3d_ versus Pre). At Post_+3d_, 14 of 17 participants improved their accuracy compared to Pre, while 3 maintained their levels (see the magenta box in suppl. Fig. 1.

Lastly, for the *mixed* group (blue histograms), there was a main effect of *test* (χ^2^=29.84; p<0.0001; Kendall’s W=0.59). Free-throw accuracy improved immediately after training for the three post-test sessions (Z>3.30, p<0.001, and r>0.80), and remained stable afterward (Z<1.54, p>0.1, and r<0.35 for comparisons among post-tests). Free-throw accuracy increased significantly immediately after training (+16.5%, Post vs Pre, p<0.001), improved slightly 24 hours later (+2.9%, Post_+1d_ vs Post, p=0.04), and then stayed stable (+0.6%, Post_+3d_ vs Post_+1d_, p=0.39). The total gain in free-throw accuracy was 20% (p<0.001; Post_+3d_ vs Pre). At Post_+3d_, 16 of 17 participants improved their accuracy compared to Pre, while one maintained it (see the blue box in suppl. Fig. 1).

### 2. Multiple Regression Analysis

As a supplementary analysis, multiple regression analyses were conducted to examine whether the SPSRQ sensitivity and reward scores predicted changes in free-throw performance, while controlling for baseline performance (Pre) and group. As in the main text, we performed a first multiple regression excluding participants who responded to the questionnaire after training (n=54), and a second one including all participants (n=68). The results were similar in both cases. We presented hereafter those including 54 participants.

For immediate improvement (Post–Pre gain; see Fig. 3A in the main text), the regression analysis revealed a significant model, F(3,50) = 11.62, p < 0.001, explaining 41.1% of the variance (R^2^ = 0.41). Baseline performance significantly predicted improvement (B = −0.63, SE = 0.14, t = −4.40, p < 0.001), and group also had a significant effect (B = 2.90, SE = 1.17, t = 2.48, p = 0.017). Punishment sensitivity was significantly associated with improvement (B = 0.95, SE = 0.43, t = 2.18, p = 0.034), suggesting that individuals with higher punishment sensitivity tended to show larger immediate gains. For offline gains (Post_+3d_–Post gain; see Fig. 3C and 3D in the main text), the regression model including reward sensitivity was significant, F(3,50) = 4.17, p = 0.01, explaining 20.0% of the variance (R^2^ = .20). Reward sensitivity significantly predicted performance change (B = 1.10, SE = 0.36, t = 3.06, p = 0.004), whereas the effects of group (p = 0.06) and baseline performance (p = 0.15) were not significant. In the model including punishment sensitivity, the overall regression was also significant, F(3,50) = 5.04, p = 0.004, explaining 23.2% of the variance (R^2^ = .23). Punishment sensitivity significantly predicted performance change (B = −1.94, SE = 0.52, t = −3.43, p = 0.001), whereas group (p = 0.81) and baseline performance (p = 0.22) were not significant predictors. Together, these results indicate that individual differences in reward and punishment sensitivity are associated with changes in performance during the offline period, independently of baseline ability and group.

